# Microbial bioprospecting for lignocellulose degradation at a unique Greek environment

**DOI:** 10.1101/2020.08.29.273201

**Authors:** Daphne N. Georgiadou, Pavlos Avramidis, Efstathia Ioannou, Dimitris G. Hatzinikolaou

## Abstract

Bacterial systems have gained wide attention for depolymerization of lignocellulosic biomass, due to their high functional diversity and adaptability. To achieve the full microbial exploitation of lignocellulosic residues and the cost-effective production of bioproducts within a biorefinery, multiple metabolic pathways and enzymes of various specificities are required. In this work, highly diverse aerobic, mesophilic bacteria enriched from Keri Lake, a pristine marsh of increased biomass degradation and natural underground oil leaks, were explored for their metabolic versatility and enzymatic potential towards lignocellulosic substrates. A wide diversity of *Pseudomonas* species were obtained from enrichment cultures where organosolv lignin served as the sole carbon and energy source and were able to assimilate a range of lignin-associated aromatic compounds. Highly complex bacterial consortia were also enriched in cultures with xylan or carboxymethyl cellulose as sole carbon sources, belonging to Actinobacteria, Proteobacteria, Bacilli, Sphingobacteriia, and Flavobacteria. Numerous individual isolates could target diverse structural lignocellulose polysaccharides by expressing hydrolytic activities on crystalline or amorphous cellulose and xylan. Specific isolates showed increased potential for growth in lignin hydrolysates prepared from alkali pretreated agricultural wastes. The results suggest that Keri isolates represent a pool of effective lignocellulose degraders with significant potential for industrial applications in a lignocellulose biorefinery.

## 1. Introduction

In the last decades, the fast-growing human population and the consequent accelerating consumption of Earth’s resources have raised global concern over our planet’s material, energy and environmental sustainability (Ferreira 2017). On the other hand, petroleum’s availability is depleting while the dependence of the transport and chemicals sector on fossil fuels remains high (Cherubini 2010). These circumstances necessitate the use of alternative sources for the production of fuels and other industrial chemicals. The biorefinery concept, which is based on the sustainable conversion of renewable biomass into marketable products, was proposed as a potential solution to help moderate the harmful consequences of the growing demand for fuels, energy, and chemicals (Kawaguchi et al. 2016).

Lignocellulose, which is the major structural component of plant cell walls, is the most abundant and fully renewable resource of reduced biomass in the biosphere (B Fisher and S Fong 2014). High amounts of lignocellulosic waste are created annually, through agricultural practice, forestry, industrial processes such as paper and pulp, textile and timber industries or breweries, and can serve as a raw material for the production of biofuels, bioplastics, and other platform chemicals, within a biorefinery (Sharma, Xu, and Qin 2019). Lignocellulose is mainly composed of a mixture of carbohydrate polymers such as cellulose and hemicelluloses, interconnected through the aromatic heteropolymer of lignin. Within cellulose microfibrils, regions of highly ordered structure create a highly organized and crystalline form with increased recalcitrance (Lee 1997). Moreover, the heterogeneity of hemicelluloses and the highly diverse structure of the lignin molecule create a substrate with increased resistance to chemical and enzymatic degradation.

Within a biorefinery, the conversion of lignocellulose into biofuels and bioproducts involves three main steps. The first step involves the delignification of plant material and the disruption of cellulose’s crystal structure, aiming to enhance the accessibility of hydrolytic enzymes to glucan and xylan (Alvira et al. 2010). Typically, it is achieved either by mechanical, physical, or chemical pretreatment processes of plant biomass. After pretreatment, hydrolytic enzymes are used to depolymerize cellulose and hemicelluloses into simple sugars and in the final step, sugars are fermented and converted to ethanol or other fuels by microorganisms.

Currently, the main enzymes used for lignocellulose hydrolysis are derived from fungi. However, their relatively high production cost, resulting from the insufficient quantities produced accounts for 20-40% of the total cost for cellulosic ethanol production (Naresh Kumar et al. 2019). The use of bacterial enzymes for the hydrolysis of polysaccharides could be more advantageous due to the wider availability and higher efficiency of heterologous expression systems for prokaryotic genes (W. Li et al. 2008). Also, bacteria display higher tolerance and stability against harsh process conditions and potential inhibitors formed during the conversion process of lignocellulose. Particularly, bacteria and their enzymes are active in neutral or alkaline pH, a beneficial trait for growth or activity in unneutralized lignin streams from alkaline pretreatments (Mathews, Pawlak, and Grunden 2015). Microbial degradation of pretreated lignin can detoxify or even transform lignin effluents into targeted products. Residual lignin is mostly underutilized since only a small amount of the lignin fraction (2%) is used for the generation of fine chemicals (Laurichesse and Avérous 2014). Chemical procedures implemented for lignin fragmentation and transformation into high-value products usually result in the formation of multiple diverse products, which necessitate an extensive separation process. In contrast, bacterial-based systems of lignin metabolism are gaining growing attention, owing to the bacteria’s ability to funnel diverse lignin-derived molecules into readily purified, high added-value products, such as vanillin, polyhydroxyalkanoates (PHA), lipids and pyruvate (Linger et al. 2014; Xu et al. 2019).

Within the above framework, our objective is to detect novel lignocellulose degrading bacteria and explore their metabolic and enzymatic potential, with the ultimate aim to identify high-efficient biocatalysts that can be employed in a lignocellulose degrading industrial bioprocess. Our efforts were focused at Keri Lake, on the island of Zakynthos, western Greece. The area represents a unique environment, dominated by a marsh mainly composed of reeds, with increased biomass degradation, where, in parallel, asphalt springs release crude oil for more than 2.500 years. Previously reported hydrocarbon analyses of surface oil samples from the study area showed an increased content in aromatic hydrocarbons (Palacas et al. 1986; Panagiotaras et al. 2012). Our hypothesis is that these characteristics create a unique habitat where indigenous microbial populations harbor specialized degradative capabilities towards cellulose and hemicelluloses, resulting from decaying biomass, and towards aromatic substances present in lignin or aromatic hydrocarbons. To test our hypothesis and isolate bacteria able to degrade lignocellulose-derived components, we used an enrichment strategy based on organosolv lignin, birchwood xylan, and amorphous cellulose, as sole carbon and energy sources. To further examine the metabolic and enzymatic potential of the isolates, single strains were screened on a wide range of cellulosic, hemicellulosic, and lignin substrates of interest.

## 2. Materials and Methods

### 2.1 Collection of soil samples

Keri Lake is situated in the southern part of Zakynthos Island, in western Greece, with its surface lying 1 m above the sea level. Nowadays, this coastal fen is characterized by increased plant biomass degradation along with natural oil seeps at several sites. The vegetation of the area is dominated by the marsh reed *Phragmites australis*, and various peat-forming plants, that form a five-meter thick peat layer below the fen (Avramidis et al. 2017). Surface soil samples (5-10 cm depth) were collected from five uniform sites of Keri Lake, exposed to oil, during a sampling campaign conducted in October 2013 (Michas et al. 2017). Soil samples were kept in sterile containers, placed in a portable fridge (4-8 °C), and transferred within 24 h in the lab. Soil samples were stored at 4 °C until further processing (maximum storage time in the lab before the commencement of enrichments was 48 h).

### 2.2 Isolation of bacterial strains from soil samples

In order to isolate aerobic lignocellulolytic bacteria, enrichment cultures were performed by suspending 2 g of each soil sample in 20 mL Minimal Salt Medium (MSM), supplemented with either 1% (w/v) carboxymethyl cellulose (CMC, Sigma - Aldrich), 1% (w/v) birchwood xylan (Sigma - Aldrich) or 1% (w/v) organosolv lignin (Sigma - Aldrich, CAS No. 8068-03-9) as sole carbon sources. NaCl concentration was adjusted to match that of the coastal waters of Keri area. MSM consisted of: (g L^−1^) NaH_2_PO_4_.H_2_O 3.8, NaH_2_PO_4_.7H_2_O 3.0, NH4Cl 0.8, NaSO_4_ 0.23, NaCl 8.5, KCl 0.5, MgCl_2_.6H_2_O 0.32, CaCl_2_.2H_2_O 0.03, supplemented with trace elements and vitamins. The initial pH was adjusted to 6.5. Cultures were incubated at 30 °C under continuous shaking at 180 rpm. Subsequent transfers were carried out (1 mL into 20 mL of fresh medium) at 5 days intervals. At the end of a 30 days enrichment period, 100 μL from suitable dilutions of each final enrichment flask were spread on agar plates containing MSM supplemented with 1% (w/v) of the corresponding carbon source, and plates were incubated at 30 °C under aerobic conditions. Bacterial colonies were selected based on their morphology and color and were repeatedly streaked and purified on nutrient agar plates.

### 2.3 Identification and phylogenetic analysis of isolated strains

Isolation of genomic DNA was carried out from overnight grown pure bacterial cultures in nutrient broth, by using the CTAB method, following the Joint Genome Institute bacterial genomic DNA isolation protocol (William, Feil, and Copeland 2012). The purified DNA was used as a template for amplification of the full-length 16S rRNA gene with Phusion DNA Polymerase, using the forward primer 27F (5'-AGAGTTT-GATCMTGGCTCAG-3') and reverse primer 1492R (5'-TACGGYTACCTTGTTACGACTT-3'). The temperature profile consisted of 98 °C for 1 min, followed by 35 cycles of denaturation at 98 °C for 10 sec, annealing at 59.5 °C for 30 sec and extension at 72 °C for 50 sec. The 1465 bp PCR products were gel-purified using a Macherey - Nagel Gel Extraction kit and sequenced. The obtained sequences were compared to two different databases, Genbank database (16S ribosomal RNA database), by using the BLASTn algorithm (http://blast.ncbi.nlm.nih.gov/Blast.cgi), and EzBio-Cloud database (http://www.ezbiocloud.net/) (Yoon et al. 2017). Multiple sequence alignments of the isolates’ 16S rRNA genes and those of the reference sequences from the GenBank database were generated using the L-INS-i algorithm of MAFFT software. The alignments served as the input for a maximum-likelihood phylogenetic tree using IQTREE version 1.6.10, with Jukes-Cantor chosen as a best-fit model according to BIC. The confidence values of branches were generated using bootstrap analysis based on 1000 iterations.

### 2.4 Screening for glycoside hydrolase activities on solid media

All isolates were screened for hydrolytic activities on agar plates containing MSM supplemented with either 1% (w/v) CMC, 1% (w/v) Avicel microcrystalline cellulose (Sigma - Aldrich) or 1% (w/v) birchwood xylan. A cell suspension of each isolate in Ringer solution was spread on each plate and growth was evaluated after a 5-10 day incubation at 30 °C. Each experiment was conducted in duplicates. Once the growth of the bacterial cells was sufficient, agar plates were overlaid with a Congo Red solution (Sigma - Aldrich), of 1 mg mL^−1^ concentration, and stained for 15 minutes. The Congo Red solution was then replaced by a 1 M NaCl rinsing solution for 15 minutes. The appearance of a clearance zone around the bacterial colonies indicated hydrolysis of the substrate and thus cellulolytic or xylanolytic activity (Teather and Wood 1982).

### 2.5 Preparation of lignin hydrolysates by alkaline pretreatment of plant biomass

Agricultural residues from corn stover and wheat straw were provided by a local producer in the province of Phthiotis, central Greece. Each raw material, consisting of stem and leaves, was air-dried and subjected to mechanical treatment, involving homogenization in a mixer and grinding through a sieve-mill (0.7 mm). Subsequently, an aqueous solution (10% w/v) of the milled material was heated at 90 °C for 30 minutes, in order to remove impurities, followed by a thorough washing on a sieve (160 μm) using tap and distilled water. The washed solid material was dried at 50 °C, until constant weight. 20 g of each dried material was further dried in a moisture balance (Kett, FD600) while their moisture content was determined. Alkaline (soda) pretreatment was carried out by adding 0.1 g NaOH per gram of dry biomass, in an aqueous solution, at a solid-liquid mass ratio of 1:6. The slurry was heated at 95 °C for 3 hours, under continuous stirring. Finally, the solution was filtered through a 160 μm sieve to separate the black liquor of solubilized lignin from the solid residues of plant material. The final alkali solubilized lignin was subjected to the following analysis: Total carbon (TC), total inorganic carbon (TIC) and total nitrogen (TN) content were simultaneously determined, applying the high-temperature (720°C) catalytic combustion (Pt/Al2O3) oxidation method (Bekiari and Avramidis 2014), using a Shimadzu TOC analyzer (TOC-VCSH) coupled to a chemiluminescence detector (TNM-1 TN unit). Reducing sugars content was measured by the DNS method (Miller 1959), while total dissolved solids were determined by evaporation of the hydrolysate solution. For ^1^H NMR spectroscopy an aqueous solution of each lignin hydrolysate containing around 0.5 g L^−1^ Total Organic Carbon, was centrifuged and the supernatant was concentrated in a rotary evaporator and dissolved in deuterated water. 1H NMR spectra were recorded on a Bruker DRX 400 MHz spectrometer, at 300 K. Hydrolysates were stored at −20 °C until use.

### 2.6 Growth studies on lignin substrates and selected aromatic carbon sources

Growth on liquid media containing lignin substrates or selected aromatic compounds as sole carbon sources was evaluated for isolates from lignin enrichment cultures, and isolates belonging to the classes of α-Proteobacteria and Actinobacteria. Bacterial strains were inoculated into nutrient agar plates and grown at 30 °C. Plates were flooded with Ringer solution and cells were harvested by scraping. Cell suspensions were used for the inoculation of 5 mL liquid cultures of MSM supplemented with one of the following carbon sources, in the corresponding concentration of Total Organic Carbon (g L^−1^): corn stover lignin hydrolysate (0.5), wheat straw lignin hydrolysate (0.5), (both prepared as described above), ferulic acid (1.0) (Acros), caffeic acid (1.0) (Extrasynthese), vanillic acid (1.0) (Acros), syringic acid (1.0) (Acros), biphenyl (0.9) (Merck) and guaiacylglycerol-betaguaiacyl ether (GGE) (0.6) (TCI). No further data about the isomer composition of GGE were available. Kraft lignin (Sigma – Aldrich, CAS Number 8068-05-1) was added at a concentration of 0.1 % (w/v). Lignin containing media were autoclaved, whereas, media with mono- and dimeric aromatic compounds were filter sterilized. Each final medium was supplemented with trace elements and vitamins, and pH was adjusted to 6.5. The inoculum volume was adjusted so as to achieve a 0.2 initial optical density in the culture. Three replications were conducted for each carbon source and inoculated MSM basal medium with no carbon source added, was implemented as control culture. Cultures were incubated at 30 °C on a rotary shaker at 180 rpm for 14 days and samples were withdrawn for determining the optical density in a microplate reader (600 nm). Strain *Pseudomonas putida* KT2440 was used as a model strain during the growth tests.

## 3. Results

### 3.1 Identification and phylogenetic analysis of isolated bacterial strains

Aerobic, mesophilic bacteria from five uniformly distributed soil samples of Keri Lake (Suppl. Figure 1), were enriched in liquid cultures containing either organosolv lignin, birchwood xylan or amorphous cellulose as sole carbon and energy sources. A total of 63 colonies were isolated from enrichment cultures of all five soil samples, named ZKA1 to ZKA63. All isolates were deposited at Athens University Bacterial & Archaea Culture Collection (ATHUBA) under the Accession numbers ATHUBa1401 to ATHUBa1463 (http://m-biotech.biol.uoa.gr/ATHUBintex.html). Among them, 19 were isolated from lignin, 23 from xylan, and 21 from cellulose enrichment cultures. Solid media were dominated by bacteria while fungal colonies were scarcely observed. Characterization of the 16S rRNA gene of all the isolates generated 24 different genera. All isolates shared >97% identity with their closest counterparts (Suppl. Table 1), and their 16S rRNA sequences were deposited in GenBank database (Accession numbers: MT683172 to MT683234). 16S rRNA gene sequences of the isolates and their related strains were used to generate a maximum-likelihood phylogenetic tree (Figure 1).

**Figure 1.**
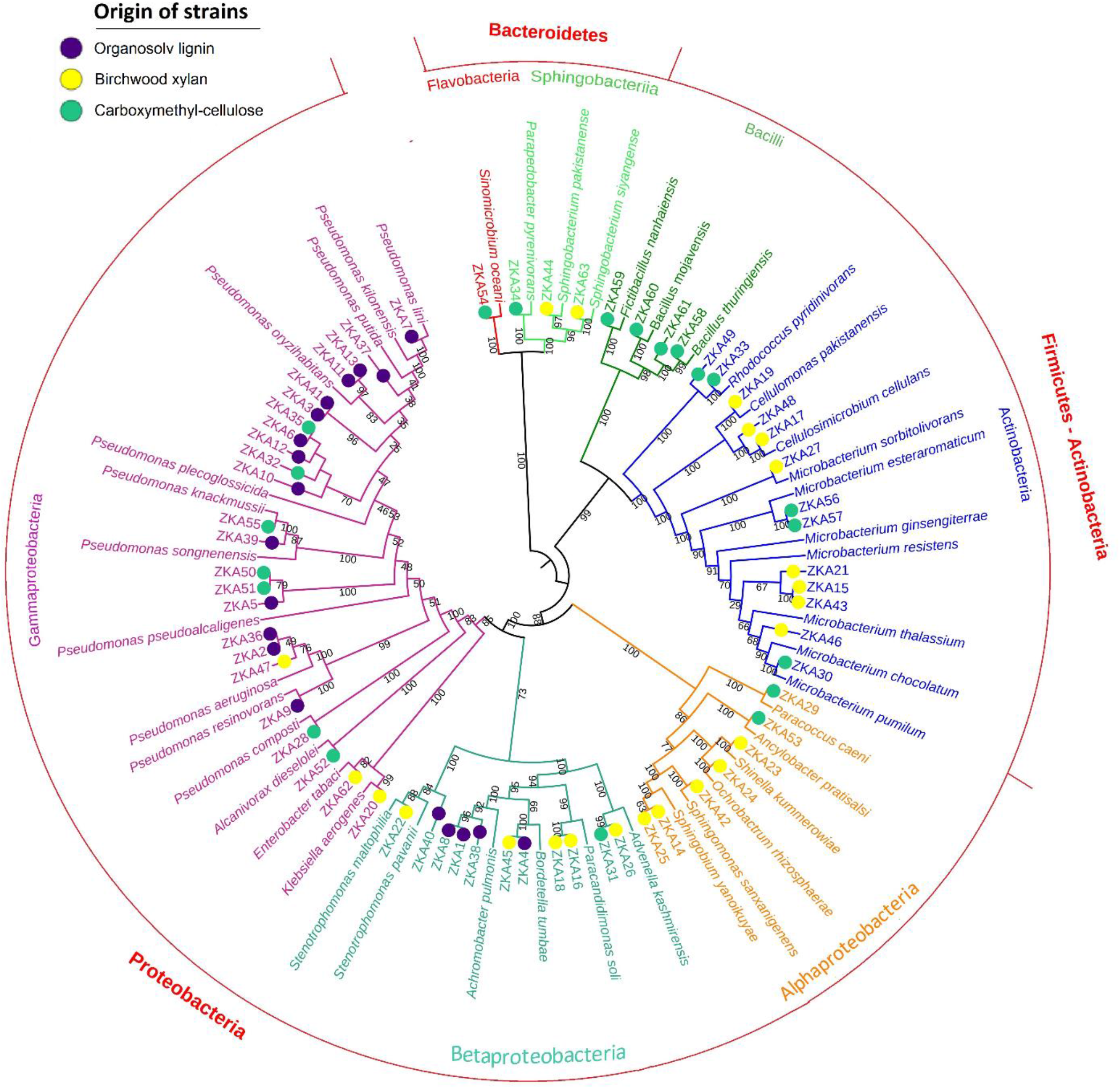
Phylogenetic tree from maximum likelihood analysis of 16S rRNA gene of Keri Lake isolates and their most similar GenBank sequences. Distance matrices were calculated by the Jukes-Cantor model. Bootstrap values expressed as percentages of 1000 replications are shown at the branch points. Colored circles indicate the origin of the bacterial strains according to the sole carbon source used in the corresponding enrichment culture; purple: organosolv lignin isolates, yellow: birchwood xylan isolates, green: carboxymethylcellulose isolates.

Sequences were grouped into 3 main clusters, with high bootstrap values of branching points (>88%), including the following groups: Bacteroidetes phylum, Firmicutes-Actinobacteria phyla, and Proteobacteria phylum. Bacteroidetes phylum was differentiated into two subclusters of Flavobacteria and Sphingobacteria classes. Firmicutes-Actinobacteria cluster included two subclusters with strains representative of the Actinobacteria and Bacilli classes (bootstrap values 100%). The cluster of the Proteobacteria phylum comprised two subclusters of the gamma-beta (γ-β) and alphaproteobacteria classes, indicated by high bootstrap values (100%). The γ-β proteobacteria cluster consisted of two branches, with representatives from γ- and β-proteobacteria classes, however, the bootstrap values of the branching points were slightly lower (85% and 73% respectively). Strains of the γ-class primarily belonged to *Pseudomonas* genus, though the sequence analysis could not reliably distinguish their species. This fact is consistent with previous studies, reporting that phylogenetic affiliation of pseudomonads based on 16S rRNA gene alone can generate misclassifications (Moore et al. 1996). Thus, further analysis of more housekeeping genes is required to correctly assign these strains. Two isolates, ZKA22 and ZKA40, though identified as *Stenotrophomonas* species, clustered in the class of β-proteobacteria. Similarly, xanthomonads group, from which some strains were later re-classified as stenotrophomonads, was previously reported to cluster either to beta- or gamma-proteobacteria, depending on the treeing algorithm and the number of rDNA sequences included (Falkow et al. 2006). Strains ZKA1, ZKA2, ZKA8, ZKA22, ZKA36, ZKA38 and ZKA47 were excluded from further experiments due to the opportunistic pathogenicity of their closest counter-parts, based on their 16S rRNA sequence.

### 3.2 Distribution of genera based on the enrichment carbon source

Our results revealed a distinct distribution of strains isolated from lignin enrichments mainly among gamma- and secondarily among beta-proteobacteria. Lignin enriched isolates were dominated by pseudomonads, accounting for 79 % of the total number of strains isolated from lignin cultures (Figure 2).

**Figure 2.**
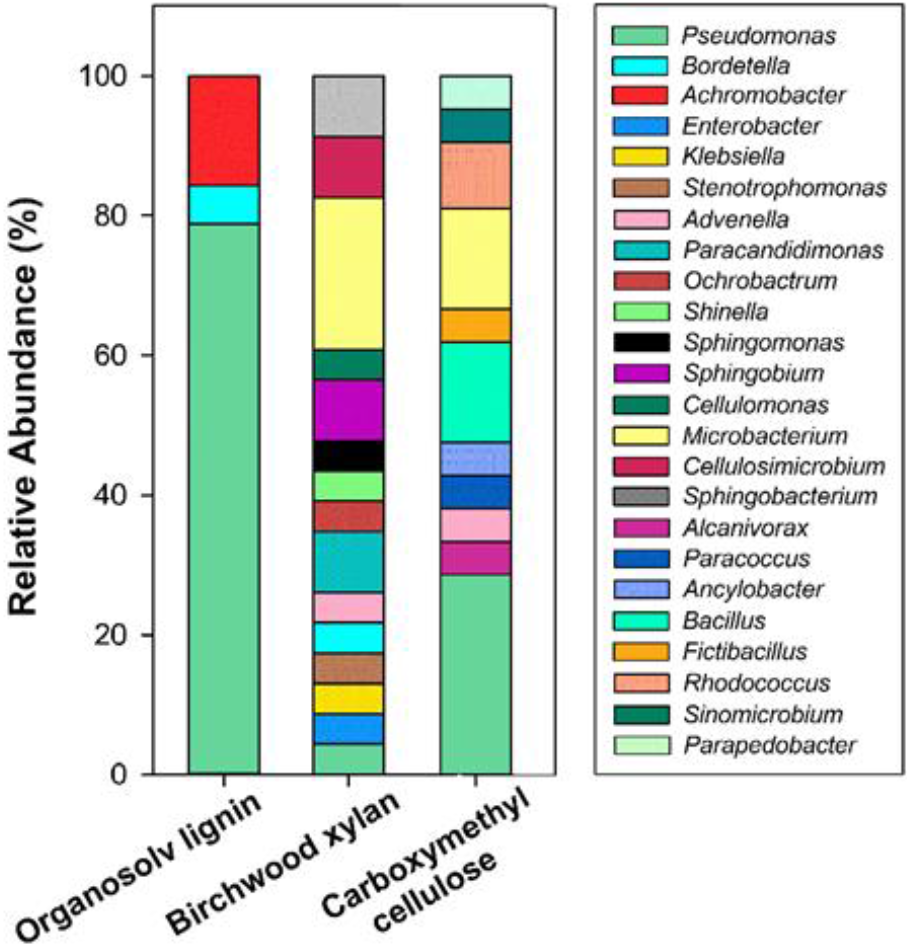
Substrate distribution of bacterial genera of Keri Lake isolates.

The rest of the isolates enriched from lignin cultures belonged to *Achromobacter* spp. (16 %) and *Bordetella* spp. (5 %). A highly complex bacterial diversity was detected among strains enriched in xylan and CMC cultures. Strains belonging to the class of Actinobacteria, alpha-Proteobacteria, Bacilli, Sphingobacteriia, and Flavobacteria were exclusively encountered in these substrates, accompanied by members of the gamma- and beta-Proteobacteria. *Microbacterium* was the most representative genus in xylan enrichment cultures, (22% relative abundance), followed by *Sphingobacterium*, *Cellulosimicrobium*, *Paracandidimonas,* and *Sphingobium* (9 % each). Strains of the genera *Pseudomonas*, *Enterobacter*, *Klebsiella*, *Stenotrophomonas*, *Bordetella*, *Advenella*, *Ochrobactrum*, *Shinella*, *Sphingomonas* and *Cellulomonas* were equally abundant (4 %), among xylan isolates. Strains enriched in CMC cultures were dominated by *Pseudomonas* (29 %), *Bacillus* (14 %), *Microbacterium* (14 %), and *Rhodococcus* (10 %) species. Members of the genera *Alcanivorax*, *Advenella*, *Paracoccus*, *Ancylobacter*, *Fictibacillus*, *Sinomicrobium,* and *Parapedobacter* were also recovered, each genus equally representing 5 % of the total number of strains enriched in this medium.

### 3.3 Screening for glycoside hydrolase activity on solid media

To further assess the capability of the isolates to express cellulolytic and xylanolytic activities, all strains were subsequently grown on plates with CMC amorphous cellulose, Avicel microcrystalline cellulose or xylan from birchwood, as sole carbon sources, and were subjected to the Congo red assay for qualitative observation of hydrolase activity. In most cases, three to five days were required for the strains to grow on the substrates and express the corresponding glycoside hydrolases (GH) activity. Almost half of the isolates (43%), regardless of their origin in enrichment cultures, could sufficiently grow on all three carbon sources and produced hydrolysis zones after staining with Congo red (Figure 3). Within the phylum level, Firmicutes possessed the highest percentage (75%) of the hydrolytic strains for all three substrates used, followed by Actinobacteria (53%), Bacteroidetes (50%), and finally representatives of Proteobacteria (34%). To compare our findings with the number of glycoside hydrolases involved in cellulose and xylan deconstruction of each genus, we also performed an analysis on the predicted number of GHs, in the available genomes of Carbohydrate Active Enzymes database (CAZy) (Supplementary Figure 2) (Lombard et al. 2014).

**Figure 3.**
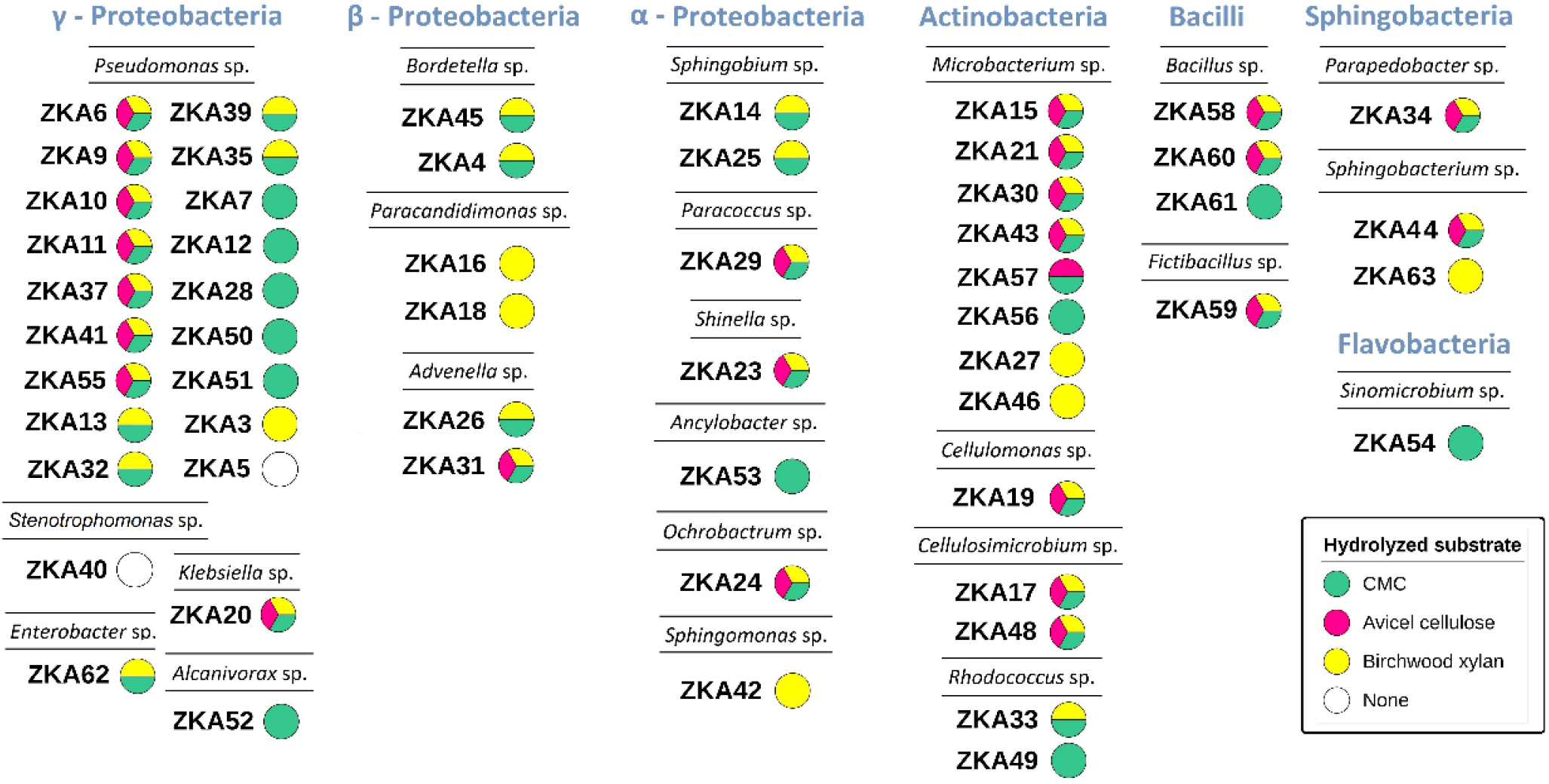
Glycoside hydrolase activity of Keri Lale isolates. Colored circles indicate the presence of hydrolysis zones on agar plates with amorphous CMC cellulose (green color), Avicel microcrystalline cellulose (red color) or birchwood xylan (yellow color) as sole carbon sources. Uncolored circles indicate lack of ability to grow on either of these substrates.

Strains *Cellulomonas* sp. ZKA19, *Cellulosimicrobium* sp. ZKA17 and ZKA48, *Sphingobacterium* sp. ZKA44, *Klebsiella* sp. ZKA20, *Microbacterium* sp. ZKA15, ZKA21, ZKA30 and ZKA43, and *Bacillus* sp. ZKA58 and ZKA60 showed the ability to deconstruct all three substrates tested. According to CAZy database, representatives of these genera contain a high diversity of GH genes in their genomes. Interestingly, the broad substrate hydrolyzing phenotype was also observed in strains ZKA6, ZKA9, ZKA10, ZKA11, ZKA37, ZKA41 and ZKA55 of the genus *Pseudomonas*. Members of *Pseudomonas* genus typically contain very few hydrolase genes, if any. Still, the numbers of predicted hydrolases for this genus varied considerably in specific isolates. Strains *Advenella* sp. ZKA31, *Fictibacillus* sp. ZKA59, *Ochrobactrum* sp. ZKA24, *Paracoccus* sp. ZKA29 and *Shinella* sp. ZKA23 could also target all three substrates tested. Although there are still few sequenced genomes for most of these genera, the number of encoded GHs is relatively low.

Strains *Enterobacter* sp. ZKA62, *Rhodococcus* sp. ZKA 33 and *Sphingobium* sp. ZKA14 and ZKA25 could process both CMC and xylan, but not Avicel cellulose. These genera typically possess more than one predicted gene for diverse GH enzymes suggesting they have increased hydrolyzing capabilities. Strains *Bordetella* sp. ZKA4 and ZKA45 could process CMC and xylan too. The absence of predicted exoglucanase genes in genomes of this genus may explain the isolates’ inability to process Avicel cellulose. Strains *Alcanivorax* sp. ZKA52 and *Ancylobacter* sp. ZKA53 could only hydrolyze CMC. Although the number of sequenced genomes of these two genera is low, few genes of potential cellulases and xylanases are encoded, possibly accounting for the incapacity of the two strains to target xylan and Avicel cellulose. Strain *Parapedobacter* sp. ZKA34 could hydrolyze all substrates of this study. Strains *Paracandidimonas* sp. ZKA16 and ZKA18 could only utilize xylan. *Sinomicrobium* sp. ZKA54 could only hydrolyze CMC. These three genera had no entries in CAZy database (as of June 2020).

### 3.4 Characterization of lignin hydrolysates from alkali pretreated plant biomass

A weak aqueous solution of sodium hydroxide was used for the alkaline pretreatment of corn stover and wheat straw, in order to prepare a hydrolysate of lignin to be used as a substrate for bacterial cultures. Residual carbohydrates were almost eliminated in the resulting lignin hydrolysates, as indicated by the very low sugar content. The corn stover lignin hydrolysate (CSLH) contained 50.1 g/L dissolved solids, of which 24.1 g/L amounted to total organic carbon (TOC) and 0.41 g/L to total inorganic carbon (TIC). The wheat straw lignin hydrolysate (WSLH) contained 43.5 g/L dissolved solids of which 22.7 g/L amounted to total organic carbon and 0.47 g/L to total inorganic carbon. Total nitrogen was equal to 0.56 g/L and 0.32 g/L for CSLH and WSLH, respectively.

Spectra of both lignin hydrolysates obtained by ^1^H NMR are shown in Figure 4. A peak at 8.4 ppm may be assigned to phenolic hydroxyl groups. Signals around 6.4 and 6.9 ppm can be attributed to aromatic protons in syringylpropane (S) and guaicylpropane (G) units respectively. The signal at 6.9 ppm in both substrates is more intense than the signal at 6.5 ppm suggesting that guaiacyl units are more abundant than syringyl units. Signals around 7.4-7.6 ppm are typically assigned to aromatic protons located in positions 2 and 6 in structures containing a carbonyl Cα=O group, to aromatic protons in positions 2 and 6 of p-hydroxyphenyl (H) units conjugated with a double bond, to protons in Cα=Cβ structures, and to aromatic protons in p-coumaric and ferulic acids. A strong signal at 5.4 ppm found in CSLH, less discernible in WSLH, could be assigned to aliphatic Hα protons in dimers containing β-5 bonds, suggesting that structures like phenylcoumaran may be present in CSLH. Signals at 4.6 ppm can be attributed to aliphatic H_β_ protons of residual β-Ο-4 structures. Signals in the region 3.5-4.4 ppm can be assigned to side chain Hγ protons in arylglycerol units and methoxyl protons (−OCH_3_). Signals around 2.6 ppm can be due to benzylic protons in β-β structures, such as resinol, and signals in the region 2.0-2.3 ppm can be assigned to aromatic and aliphatic acetate groups.

**Figure 4.**
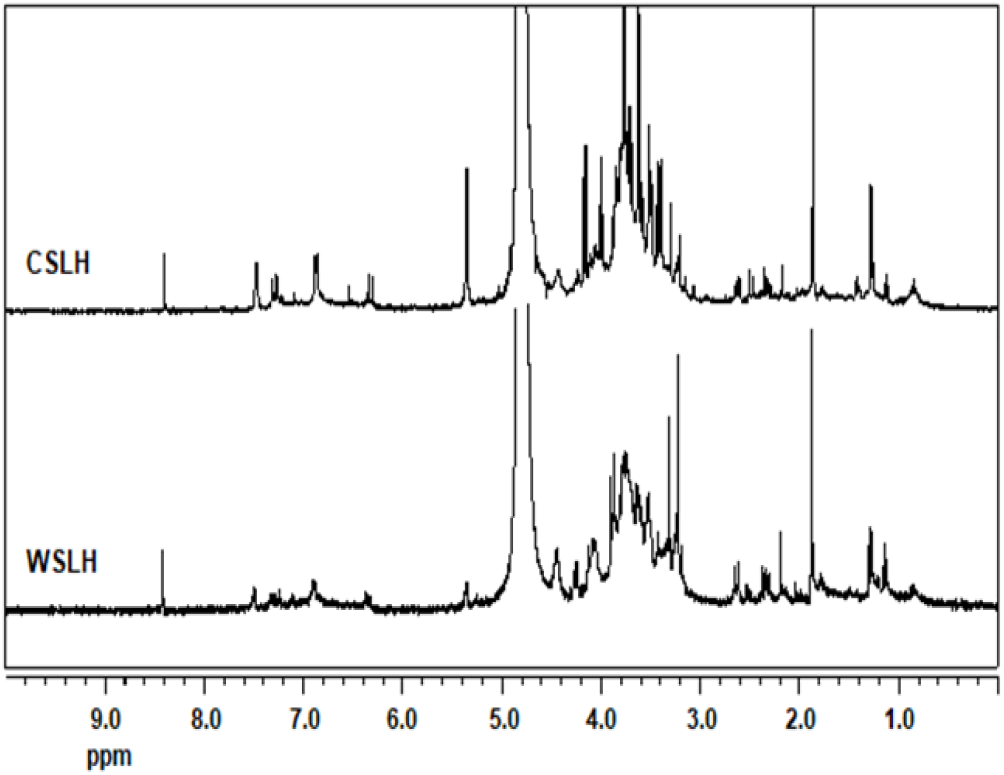
^1^H NMR spectra of lignin hydrolysates from alkali pretreated corn stover (CSLH) and (WSLH), dissolved in D_2_O and recorded on a Bruker DRX 400 MHz spectrometer, at 300 K.

### 3.5 Bacterial growth studies on different lignins

Certain strains were selected for further study of their ability to grow on mineral salt medium with lignin hydrolysates from alkali pretreated corn stover and wheat straw, or kraft lignin, as sole carbon sources. Selected strains included lignin-enriched isolates, belonging primarily to γ-Proteobacteria and secondly to β-Proteobacteria as well as strains of Actinobacteria, by reason of the numerous documented reports concerning degraders of lignin and lignin-derived compounds belonging to this class (Bugg et al. 2011). α-Proteobacteria were also included, since one of the best-characterized strains, known for its ability to degrade a plethora of lignin-derived compounds, *Sphingobium* sp. SYK-6, belongs to this class (Eiji Masai, Katayama, and Fukuda 2007). Strain *Pseudomonas putida* KT2440, known for its wide aromatic substrate – utilizing activity and its ability to grow in alkaline pretreatment liquors from corn stover (Linger et al. 2014; Belda et al. 2016; Ravi et al. 2017) was used as a control strain.

Certain microbes exhibited sufficient growth in CSLH and WSLH, while for others, growth levels were minor (Figure 5). The maximum optical density for most of the isolates that grew well in CSLH was reached within 24 or 48 hours whilst growth for the majority of them in WSLH was slower and reached its maximum after approximately 72 hours of incubation. No growth was observed in control cultures inoculated with each strain, in the absence of a lignin carbon source. Higher growth levels in CSLH were obtained by *Pseudomonas* sp. ZKA7, followed by *Bordetella* sp. ZKA4, *Shinella* sp. ZKA23, *Cellulosimicrobium* spp. ZKA17 and ZKA48, *Microbacterium* sp. ZKA21, *Rhodococcus* spp. ZKA49 and ZKA33 and *Microbacterium* sp. ZKA46. Most of these strains exhibited higher growth levels in WSLH too. Strain ZKA23 showed the highest growth rate in WSLH, followed by ZKA48, ZKA46, ZKA7, ZKA4, ZKA17 and ZKA49. *Pseudomonas* strains ZKA37, ZKA6, ZKA5, ZKA13, ZKA3, ZKA39 and *Stenotrophomonas* sp. ZKA40 also showed adequate growth in CSLH, but for some of them cell growth in WSLH was proportionally lower than others. For *Pseudomonas putida* KT2440 growth in both substrates was less significant, similarly to the rest of pseudomonads and isolates tested. Most strains were unable to grow in kraft lignin, and only *Shinella* sp. ZKA23, *Microbacterium* sp. ZKA30, *Pseudomonas* sp. ZKA3 and *Sphingobium* sp. ZKA25 displayed distinguishable but relatively low growth in this medium.

**Figure 5.**
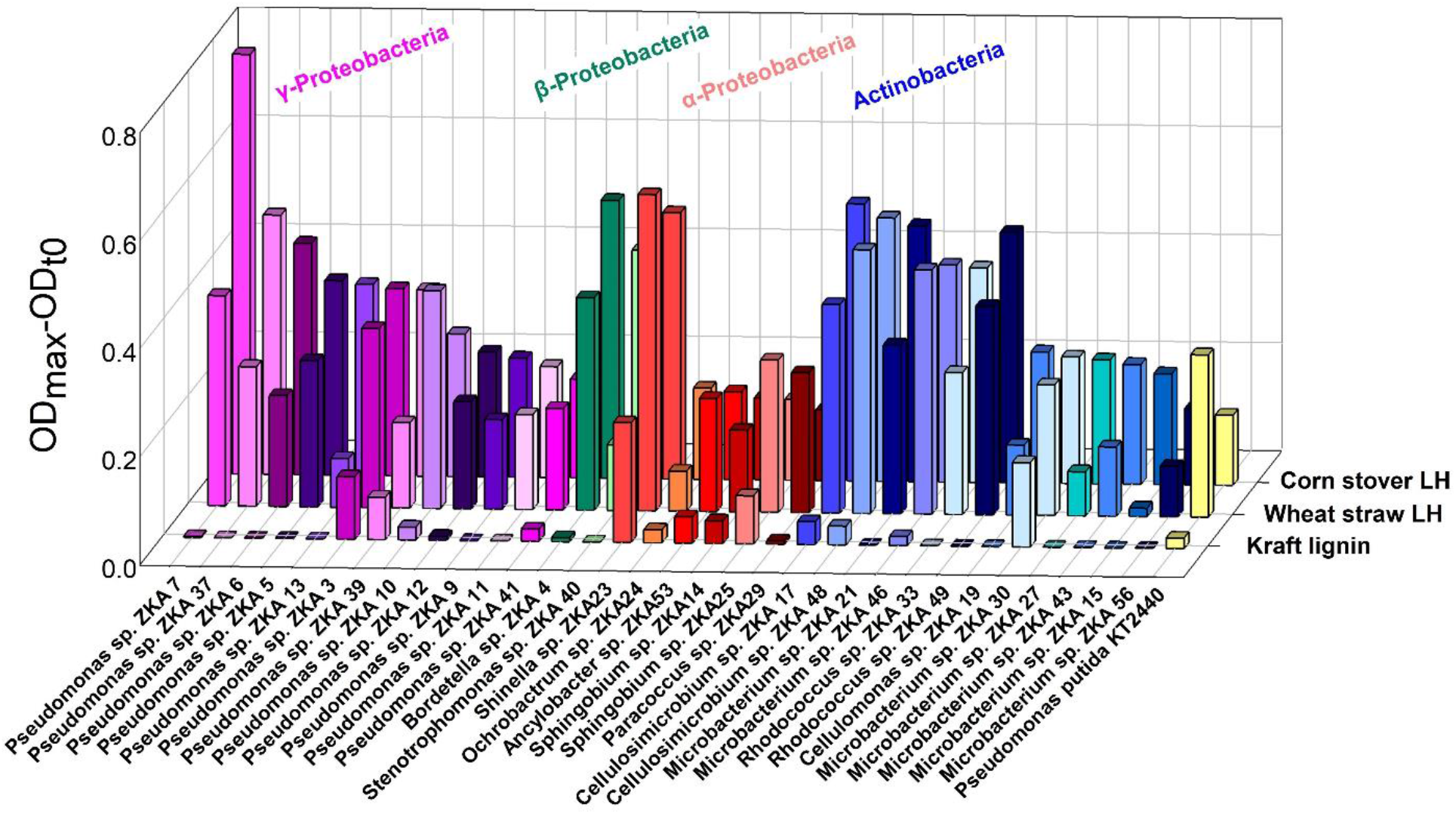
Growth of selected Keri Lake isolates in technical lignins. Isolates were grown in mineral salt medium supplemented with lignin hydrolysate (LH) from either alkali pretreated corn stover or wheat straw, or commercial kraft lignin, as sole carbon sources. Growth is expressed as difference between maximum and initial optical density.

### 3.6 Growth studies on selected aromatic carbon sources

To further elucidate their metabolic capabilities, the above-selected strains were cultivated in liquid cultures of different monoaryl or biaryl compounds as sole carbon sources. Eight out of twelve *Pseudomonas* strains showed ability to assimilate ferulic acid, which correlated well with growth in caffeic and vanillic acid, too (Table 1). The most actively growing species in these carbon sources (indicated by ++++ in Table 1) were strains ZKA3, ZKA7, ZKA10 and ZKA37. Strain *Pseudomonas putida* KT2440, used as a control strain, was also able to grow in ferulic, caffeic and vanillic acid among the substrates tested. Interestingly, *Pseudomonas* sp. ZKA12 was the only strain, amongst all strains tested, able to metabolize syringic acid under those conditions. Among isolates of Actinobacteria class, two *Rhodococcus* strains, ZKA33 and ZKA49, were able to rapidly metabolize vanillic acid and strain *Microbacterium* sp. ZKA46 was able to slowly grow on biphenyl. Alpha-proteobacterium *Ancylobacter* sp. ZKA53 was also able to grow on biphenyl. No growth was observed for the rest of the tested isolates on any carbon source.

**Table 1.**
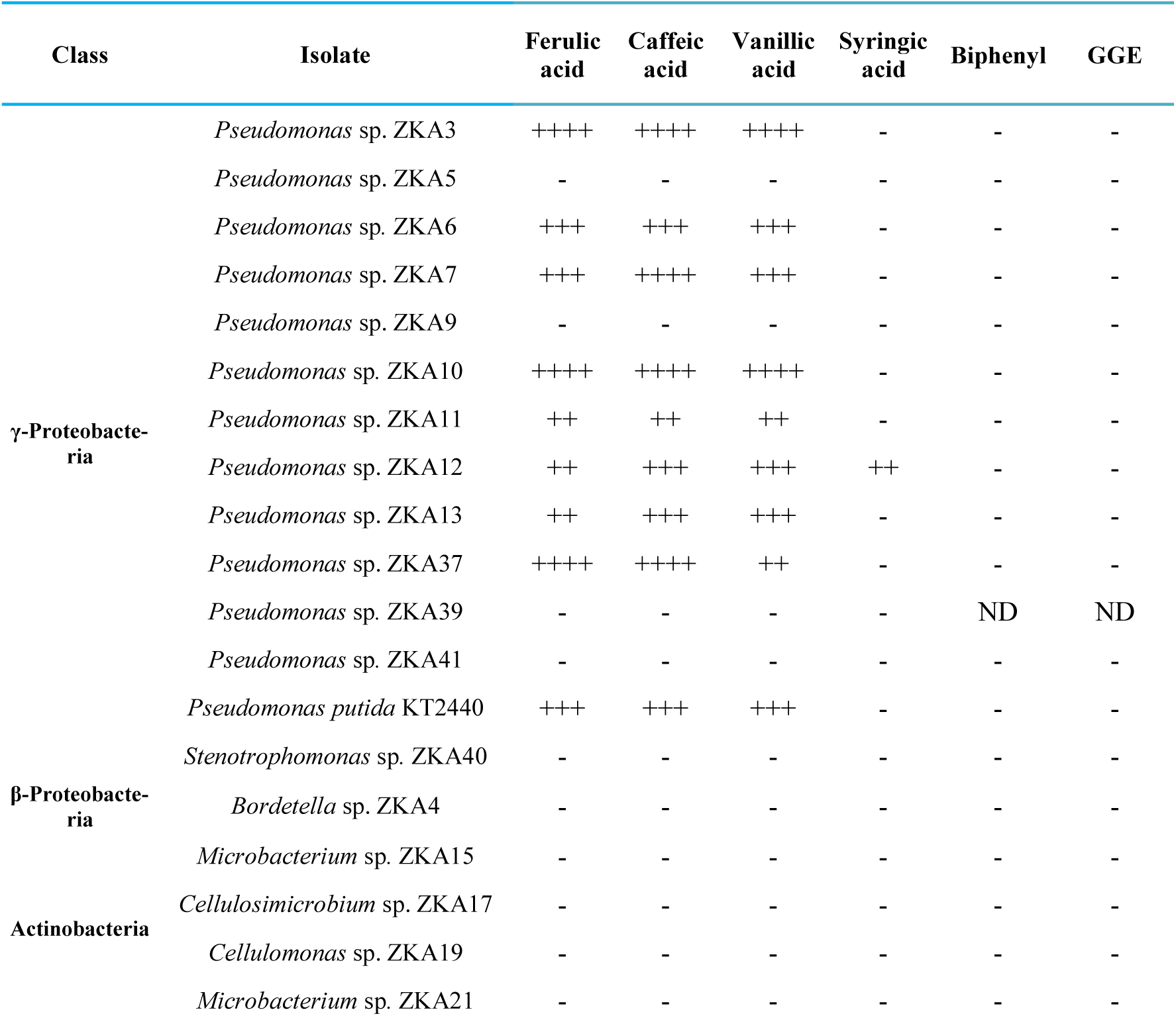
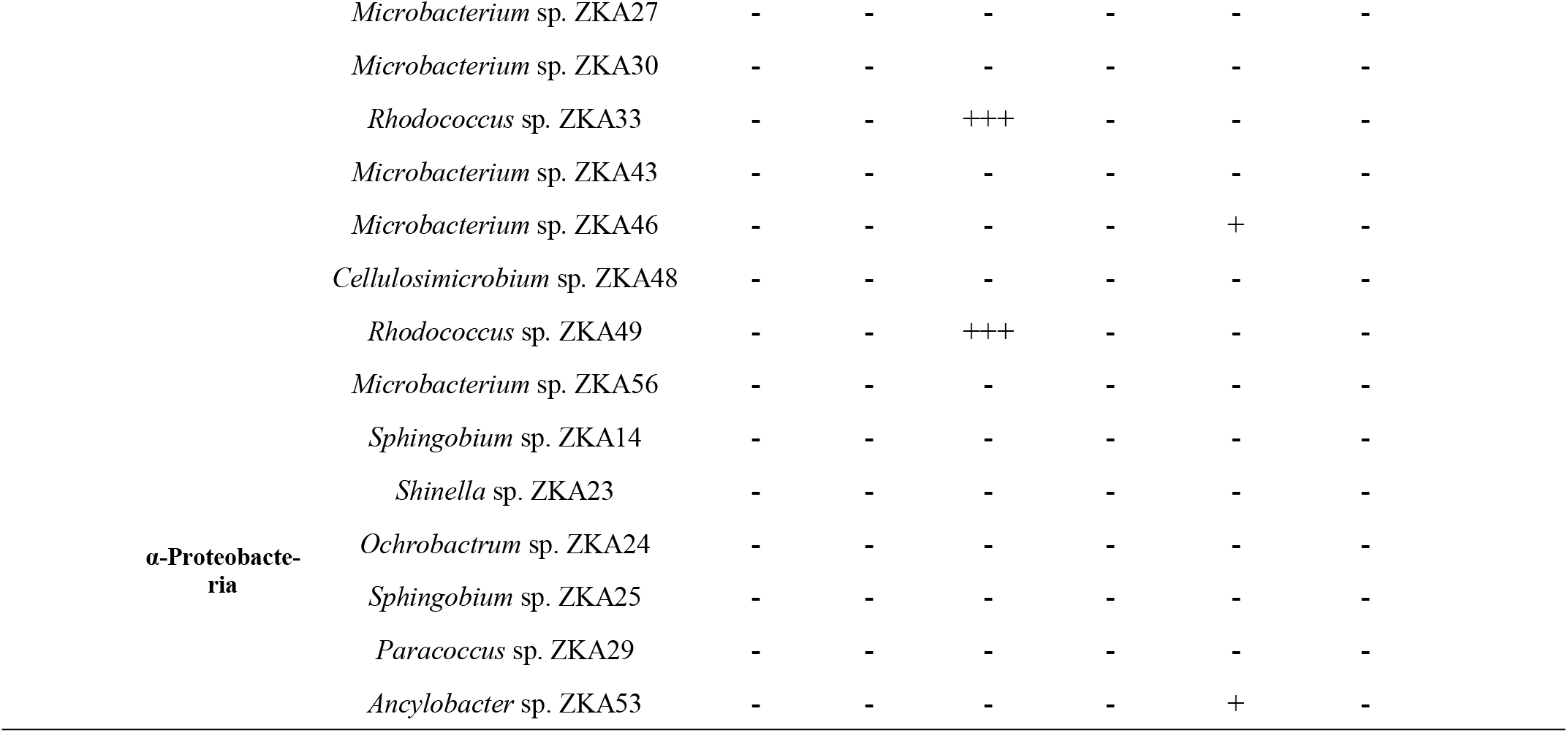
Growth of selected bacterial strains on different aromatic compounds. Isolates were grown in mineral salt medium supplemented with the corresponding compound as a sole carbon source, in a final concentration of TOC (g L^−1^): ferulic, caffeic, vanillic and syringic acid: (1.0), biphenyl (0.9), GGE (0.6). Growth is expressed as follows: (++++): OD>0.6 in 24h, (+++): OD>0.6 in 24-48h, (++): OD>0.6 in 48-96h, (+): OD>0.4 in 96-168h, (-): no growth, ND: not determined.

## 4. Discussion

Several ecosystems naturally exposed to oil are known in Western Greece, nonetheless, to our knowledge, this is the first report on the isolation of microorganisms from one such. Soil samples from Keri Lake were characterized by the presence of oil and heavily decomposed plant material. It is common for soil habitats, such as those with high phenolic content or those with increased biomass decay, to host microbes that have developed resistance and the ability to utilize the respective compounds. In fact, a metagenomic analysis on up to three-meter depth sediments from oil-exposed sites in the Keri area identified an abundance of genes involved in the anaerobic degradation of phenolic compounds (Michas et al. 2017). In our study, an enrichment approach based on three distinct lignocellulosic substrates (organosolv lignin, xylan and carboxymethylcellulose) was used for the isolation of lignocellulose degraders. Despite the limitations introduced by the enrichment technique, the number of different bacterial genera encountered was high, possibly reflecting the active rich microbial diversity of the Keri area ecosystem.

For the isolation of lignin degraders we used organosolv lignin as the sole carbon source. This type of lignin is considered relatively purer compared to other technical lignins, due to its sulfur-free structure, and lower ash and carbohydrate content. Although the effect of organosolv pretreatment on the valorization of sugars, and the use of lignin as fuel or as a precursor for chemicals have been extensively studied (Zhao, Cheng, and Liu 2009; de la Torre et al. 2013), little is known about the microbial utilization of organosolv lignin. There is only one report on the bacterial utilization of effluents obtained from organosolv pretreatment of pine, enriched with degraded oligosaccharides and lignin, by the oleaginous soil bacterium *Rhodococcus opacus* DSM 1069 (Wells, Wei, and Ragauskas 2015). In the present work, we were able to isolate several bacterial strains from organosolv lignin cultures, belonging mainly to the gamma- and secondarily to the beta-proteobacteria. Bacteria belonging to the Proteobacteria phylum are commonly isolated from lignin and lignin model compounds enrichment studies (Brink et al. 2019). Among them, many strains fall into *Pseudomonas* genus, as reported by several studies (Bandounas et al. 2011; Ravi et al. 2017; Hirose et al. 2013). Although it is difficult to discriminate between the numerically abundant strains in the natural environment and strains that display opportunistic traits in the enrichment cultures, *Pseudomonas* species are indeed widespread in soil, rhizosphere, aquatic habitats and areas polluted with manmade and natural toxic chemicals (Jiménez et al. 2010). Large sets of unique catabolic genes for a wide range of substrates, transporters, regulators and stress response systems are features accounting for the great metabolic versatility and adaptability to diverse environmental conditions of members of this genus (Dos Santos et al. 2004). Further assessment of these strains, in pure cultures or consortia, could reveal their potential for use in an industrial process of organosolv lignin biotransformation into valuable chemicals, or bioremediation of sites contaminated with industrial effluents. The use of functional microbial consortia, composed of multiple strains obtained by enrichment cultures, can be a valuable approach for the complete decomposition of lignocellulosic industrial wastes, such as lignin-rich streams. Such consortia could provide complementary enzymatic activities thereby increasing the efficiency of a waste treatment process.

In our study, the xylan and CMC enrichment groups revealed the highest diversity, including bacteria that belong to almost every major bacterial phylum. Xylan is the major component of hemicelluloses, contributing over 70% of their structure. In nature, the complete depolymerization of xylan requires the cooperative activity of enzymes with diverse specificities, collectively known as xylanases. The structure of birchwood xylan consists mainly of a β-(1,4)-linked D-xylopyranose backbone chain (86%), substituted with α-(1,2)-linked 4-Ο-methyl-D-glucuronic acid (9%) (Saha 2003). Hydrolysis of the xylan backbone is mainly accomplished by endo-β-1,4-xylanases (EC 3.2.1.8) and β-xylosidases (EC 3.2.1.37) while hydrolysis of its (1→2)-α-D-(4-*O*-methyl)glucuronosyl links is achieved by 1,2-α-glucuronidases (EC 3.2.1.131) (Shallom and Shoham 2003). Xylan decomposition is thus widespread in all natural environments with commonly reported bacteria that produce xylanases including members of genera such as *Bacillus*, *Cellulomonas*, *Micrococcus*, *Staphylococcus*, *Paenibacillus*, *Arthrobacter*, *Microbacterium*, *Rhodothermus* and *Streptomyces* (Chakdar et al. 2016). The ability to degrade amorphous cellulose, such as CMC, is also not a rare trait among bacteria. The depolymerization of this artificial soluble substrate is a prerequisite but not proof of native cellulose decomposition (Koeck et al. 2014). A large number of microorganisms inhabitants of soil, forests, insects’ and animals’ digestive system or composts have been repeatedly isolated as cellulolytic (López-Mondéjar et al. 2016). Aerobic cellulolytic bacteria include species of the genera *Cellulomonas*, *Bacillus*, *Pseudomonas*, *Streptomyces*, *Micromonospora*, *Rhizobium*, *Sphingomonas*, *Ochrobactrum*, *Arthrobacter*, *Alcaligenes* and *Serratia* (McDonald, Rooks, and McCarthy 2012).

Further evidence of the isolates’ cellulolytic and hemicellulolytic capabilities was provided by qualitative screening of pure cultures on solid media containing CMC amorphous cellulose, Avicel microcrystalline cellulose or xylan from birchwood, as sole carbon sources, stained with Congo Red. Strains belonging to genera such as *Cellulomonas*, *Cellulosimicrobium*, *Sphingobacterium*, *Klebsiella*, *Microbacterium* and *Bacillus* displayed hydrolyzing activities towards all three substrates tested. Hydrolysis of CMC is correlated with endo-1-4-β-glucanases (EC 3.2.1.4), enzymes that randomly hydrolyze internal bonds of amorphous regions of cellulose, generating new ends, and β-glucosidases (EC 3.2.1.21), enzymes that hydrolyze cellodextrins and cellobiose to glucose. Avicel is a form of insoluble micro-crystalline cellulose, with a lower molecular mass and relatively low accessibility. Hydrolysis zones on Avicel agar plates provide evidence of exoglycolytic enzyme activity. Exoglucanases (or Avicelases) act on the reducing (EC 3.2.1.176) or nonreducing (EC 3.2.1.91) chain ends of cellulose, releasing cellobiose or glucose as main products. Clearing zones around colonies on xylan agar plates treated with Congo Red, that specifically binds to β-1,4 glycosidic linkages, are indicative of endo-β-1,4-xylanases (EC 3.2.1.8) that release xylooligomers from the xylan backbone and β-xylosidases (EC 3.2.1.37) that further hydrolyze oligomers to xylose.

Cellulases and xylanases are important components of enzyme cocktails for the biorefining of lignocellulosic biomass. Many commercial cocktails have inadequate glucosidase and xylanase activities and also require a stronger activity of xylosidases, to prevent end-product inhibition (Dumon et al. 2012). Mesophilic representatives of some of the above genera have been shown to produce GHs with remarkable properties, such as a β-xylosidase from *Cellulomonas fimi* with high activity at 100°C and strong alkaline pH (Kane and French 2018). The reports on characterized hydrolases showing experimental cellulolytic or xylanolytic activity for most of the genera of this study are scarce (Supplementary Figure 2). Some isolates, belonging to *Pseudomonas*, *Advenella*, *Fictibacillus*, *Ochrobactrum*, *Paracoccus* and *Shinella* genera also showed multienzymatic activity although their relative species contain on average few GHs per genome. Though the presence of a gene does not directly indicate activity, the number and diversity of putative genes coding for GHs in bacterial genomes can provide a theoretical basis for the prediction of their ability to decompose polysaccharides (Berlemont and Martiny 2015). The results suggest that certain isolates of Keri Lake may have distinctive potential for processing plant cell-wall poly- and oligosaccharides. Other genera, such as *Parapedobacter*, *Paracandidimonas* and *Sinomicrobium* had no sequenced genomes, reflecting the low number of cultivated representatives. Quantitative estimation of the enzymes’ hydrolyzing activity and characterization of their properties will further evaluate their industrial potential.

Pure strains of organosolv lignin isolates, Actinobacteria and β-proteobacteria were further assessed for their ability to grow on mineral salt medium supplemented with lignin hydrolysates from alkali pretreated corn stover or wheat straw, or supplemented with commercial kraft lignin, as sole carbon sources. A mild alkaline pre-treatment method with NaOH was used to prepare the lignin hydrolyzates from the agricultural residues, during which the alkali labile lignin linkages, such as α- and β-aryl ethers, and glycosidic bonds in carbohydrates are disrupted, causing the dissolution and degradation of lignin and hemicelluloses with lower alkali stability (Chen et al. 2013). Typically, lignin from Gramineae plants is highly soluble in alkali, leading to high extraction yield of lignin in the hydrolysates, even with mild alkaline pretreatment conditions. ^1^H NMR analysis of lignin hydrolysates revealed the presence of typical lignin resonances, (S. Li and Lundquist 1994; Seca et al. 2000; She et al. 2010), although resonances in lignin spectra may shift depending on conditions or solvents chosen and further analysis is needed to confirm the estimated assignments of proton NMR signals. However, consistent with the assignments in the lignin ^1^H NMR spectra is the fact that grass lignins are characterized mostly as guaicyl-type and that substructures found in native corn stover and wheat straw lignin are β-Ο-4’ alkylaryl ethers, followed by lower amounts of phenylcoumarans, resinols and spirodienones. The presence of p-coumarates and ferulates in corn stover and wheat straw lignin has also been previously demonstrated (del Río et al. 2012; Sammons et al. 2013; Min et al. 2017).

Certain strains such as *Pseudomonas* sp. ZKA7, *Bordetella* sp. ZKA4, *Shinella* sp. ZKA23, *Cellulosimicrobium* spp. ZKA17 and ZKA48, *Microbacterium* sp. ZKA21, *Rhodococcus* spp. ZKA49 and ZKA33 and *Microbacterium* sp. ZKA46, were able to grow efficiently in corn stover (CSLH) and wheat straw (WSLH) lignin hydrolysates. In most cases, growth levels in WSLH were slightly lower than in CSLH. It is possible that a higher lignin content in CSLH, as indicated by the more intense peaks in ^1^H NMR spectra, supports higher growth levels of bacteria. The impact of the pretreatment conditions on lignin solubilization, in both plants, needs further investigation.

Yet, in contrast to technical lignins that are pretreated more severely, such as Kraft lignin that could not support the growth of the strains tested, alkali lignin generated with mild conditions was more compatible with microbial growth. Solid lignin from chemically, mechanically and enzymatically pretreated corn stover, subjected to base-catalyzed depolymerization under mild temperatures (<140°C) and base concentrations (1% NaOH), generated higher concentrations of monomeric and dimeric lignin-derived compounds, which were readily metabolized by certain microbial strains (Rodriguez et al. 2017). In contrast, during the kraft process, non-specific chemical reactions generate a variety of compounds, rendering Kraft lignin highly irregular (Mathews, Pawlak, and Grunden 2015). Sulfur compounds introduced into the lignin molecule during this process may act as bacterial cell growth inhibitors or have negative effects on extracellular oxidoreductases involved in the decomposition of lignin. Besides, the higher molecular weight (Mw) of Kraft lignin (Mw ∼10.000 g/mol), in comparison to the lower Mw of more fragmented lignins, such as organosolv (Mw ∼ 1.000 −2.800 g/mol) (Constant et al. 2016) may further hamper its assimilation by bacteria (Ravi et al. 2017).

To test the latter assumption, Kraft lignin was size - fractionated using gel filtration chromatography on Sephadex G-25, yielding two fractions, corresponding to high (> 5.000 g/mol) and low molecular weight kraft lignin (<5.000 g/mol). Isolates selected above were examined for their ability to grow in MSM supplemented with either fraction, in a final concentration of 0.1 (w/v) of kraft lignin. Still, almost the same growth pattern presented in Figure 5 was recorded either in low or high molecular weight fraction (data not shown). Only a small number of aerobic mesophilic bacteria have been reported to degrade Kraft lignin, some being isolated from pulp and paper industry waste where the kraft process is implemented. Some of them, also, fall into genera highlighted in this work, such as *Pseudomonas putida* NX-1, *Ochrobactrum tritici* NX-1 and *Rhodococcus jostii* RHA1 (Sainsbury et al. 2013; Xu et al. 2018). However, the chemical composition and functionality, purity and molar mass of technical lignins used are highly variable, depending on the origin of lignin and the conditions of the pretreatment process used (Ragauskas et al. 2014).

The above results feature the potential of certain isolates to proliferate in lignin effluents derived from a mild alkaline pretreatment of residual plant biomass, in a lab-scale biorefinery process. Growth of bacteria in lignin-rich waste streams, generated with low energy input, without extra carbon sources added, could reduce the overall cost of a biological treatment process. Even strains with lower growth levels in these substrates may still funnel a portion of carbon into valuable metabolites rather than allocate it into biomass. Future work will elucidate any structural changes on lignin or bioproducts formed by microorganisms of this study.

Finally, the ability of the above selected strains to assimilate lignin-associated aromatic compounds was investigated. The results show that degradation of hydroxycinnamates, such as ferulate and caffeate, is a common trait among pseudomonads but not among the rest of the bacterial strains tested. Ferulic acid bears a guaiacyl nucleus and is co-polymerized into the lignin polymer by covalently bonding to xylan side-chain residues (Karlen et al. 2016). A catabolic pathway involved in the degradation of ferulate and its precursor caffeate, found in several *Pseudomonas* strains involves funneling into vanillate, through specific enzymatic reactions that produce valuable products such as vanillin (Xu et al. 2019). Vanillin is further oxidized to vanillate which is subjected to O-demethylation to produce protocatechuate (PCA). Subsequently, PCA undergoes oxidative ring-opening, catalyzed by dioxygenases, through either an *ortho*-cleavage pathway (*β*-ketoadipate pathway, *β*-KAP) or a *meta*-cleavage pathway, generating tricarboxylic acid (TCA) cycle intermediates (Eiji Masai, Katayama, and Fukuda 2007; Kamimura et al. 2017). Useful intermediate metabolites, such as *cis*, *cis*-muconic acid, a chemical precursor for adipic acid production (Vardon et al. 2016), or acetyl-CoA, a precursor for PHA or lipid biosynthesis can be synthesized through the *β*-ketoadipate pathway (Kosa and Ragauskas 2012). Elucidation of the isolates’ metabolomes and transformation pathways of lignin-derived aromatics could enable the design of engineered strains that control the further degradation of any valuable metabolites and lead to their accumulation in bacterial cultures.

The ability of strain ZKA12 to assimilate a wider range of lignin-derived aromatics with guaiacyl nuclei, such as ferulate, caffeate and vanillate, and syringyl nuclei such as syringate, would be of significant benefit for designing a single organism with multiple metabolic pathways for lignin degradation. Syringic acid carries two methoxy groups on its aromatic ring, rendering its degradation more difficult than lignin molecules with guaiacyl or p-hydroxyphenyl structure respectively (Xu et al. 2019). This is also reflected by the small number of studies that report the catabolism of syringate by aerobic bacterial species. Some of these include the well-studied lignin degrader *Sphingomonas paucimobilis* SYK-6, and the bacterial isolates *Pandoraea norimbergensis* LD001, *Pseudomonas* sp. LD002, *Bacillus* sp. LD003, *Serratia* sp. JHT01, *Serratia liquefacien* PT01, *Pseudomonas chlororaphis* PT02, *Stenotrophomonas maltophilia* PT03 and *Oceanimonas doudoroffii* JCM21046T (Katayama et al. 1988; Bandounas et al. 2011; Numata and Morisaki 2015; Tian, Pourcher, and Peu 2016).

For few strains, such as ZKA12 and ZKA11 the ability to assimilate aromatic monomers with guaicyl and/or syringyl structure, respectively, did not accord with their relatively lower growth levels on lignin hydrolysates, while others, such as ZKA4, ZKA23, ZKA17, ZKA48 and ZKA21 could efficiently metabolize lignin hydrolysates despite their inadequacy in utilizing the aromatic monomers or dimers used in this study. Perhaps some strains cannot further depolymerize or show lower tolerance towards lignin hydrolysates or some may preferentially degrade other lignin constituents or the propanoid side chains of lignin rather than the aromatic rings.

Biphenyl occurs naturally in crude oil and its degradation proceeds through a pathway that converts biphenyl into benzoate, further metabolized by a catechol *ortho*- or *meta*-cleavage pathway. Cleavage of unsubstituted biphenyls, however, does not correlate with ability to cleave the C-C biphenyl linkage of lignin-derived compounds, as the corresponding hydrolases involved share no sequence similarity or substrate specificity (Peng et al. 1999). Some of the few bacterial strains known to utilize biphenyl as a sole carbon source belong to genera such as *Microbacterium*, *Ochrobactrum*, *Pseudomonas, Rhodococcus* and *Sphingobium* (Taylor et al. 2012; Tian, Pourcher, and Peu 2016). Our results are not consistent with these reports as only strain *Microbacterium* sp. ZKA46 was able to utilize biphenyl. However, to our knowledge, this is the first report of biphenyl utilization by an *Ancylobacter* species.

Guaiacylglycerol-β-guaiacyl ether (GGE) is a model dimer for the study of lignin degradation. The ether link of this compound constitutes the most abundant linkage of the lignin polymer. However, enzymes with Cα-dehydrogenase and β-etherase activity, involved in degradation of β-aryl ether dimers, rarely occur in nature and are mainly found in Sphingomonadaceae family or the broader class of α-proteobacteria, respectively (Kamimura et al. 2017). Still, none of the isolates of this study belonging to this class could use GGE as a sole carbon source. So far, very few bacterial strains have been reported to degrade GGE, such as *S. paucimobilis* SYK-6, *Novosphingobium* sp. MBES04, *Bacillus atrophaeus* B7, *Bacillus pumilus* C6 and *Trabulsiella* sp. IIPTG13 (E. Masai et al. 1989; Huang et al. 2013; Ohta et al. 2015; Suman et al. 2016). Notably, among them, only strains SYK-6 and IIPTG13 were able to use GGE as a sole carbon source.

## 5. Conclusions

Keri area represents a unique ecosystem in Greece, due to the long-term presence of natural asphalt springs and the decay of plant biomass, providing a natural habitat for proliferation of aromatics and polysaccharide degrading microorganisms. The isolation of highly diverse lignocellulose degraders, distributed among many different genera and the large number of strains positive for cellulase, hemicellulase and lignin-associated compounds degrading activity was proof of our initial hypothesis. Hopefully this work will lay the ground for future studies on highlighting enzymes and pathways of interest, involved in the bacterial upgrading of lignocellulosic biomass.

## Acknowledgments

This research is cofinanced by Greece and the European Union (European Social Fund - ESF) through the Operational Programme «Human Resources Development, Education and Lifelong Learning» in the context of the project “Scholarships programme for post-graduate studies - 2^nd^ Study Cycle” (MIS-5003404), implemented by the State Scholarships Foundation (IKY).

## Author contributions

DGH conceived the research idea and supervised the research project. DGH and PA participated in the sampling campaign and performed soil sampling. DNG and DGH designed the experiments. DNG performed the experiments, PA performed elemental analysis of lignin hydrolysates and EI performed _1_H NMR spectroscopy and contributed to the interpretation of _1_H NMR spectra. DNG analyzed the results and wrote the manuscript draft. All authors provided critical revision of the manuscript. DH edited the final version of the manuscript.

