## Supplemental Figure1 for "Microbial bioprospecting for lignocellulose degradation at a unique Greek environment" (from another version of this preprint: version 1, 2020-08-29)

**Figure S1**

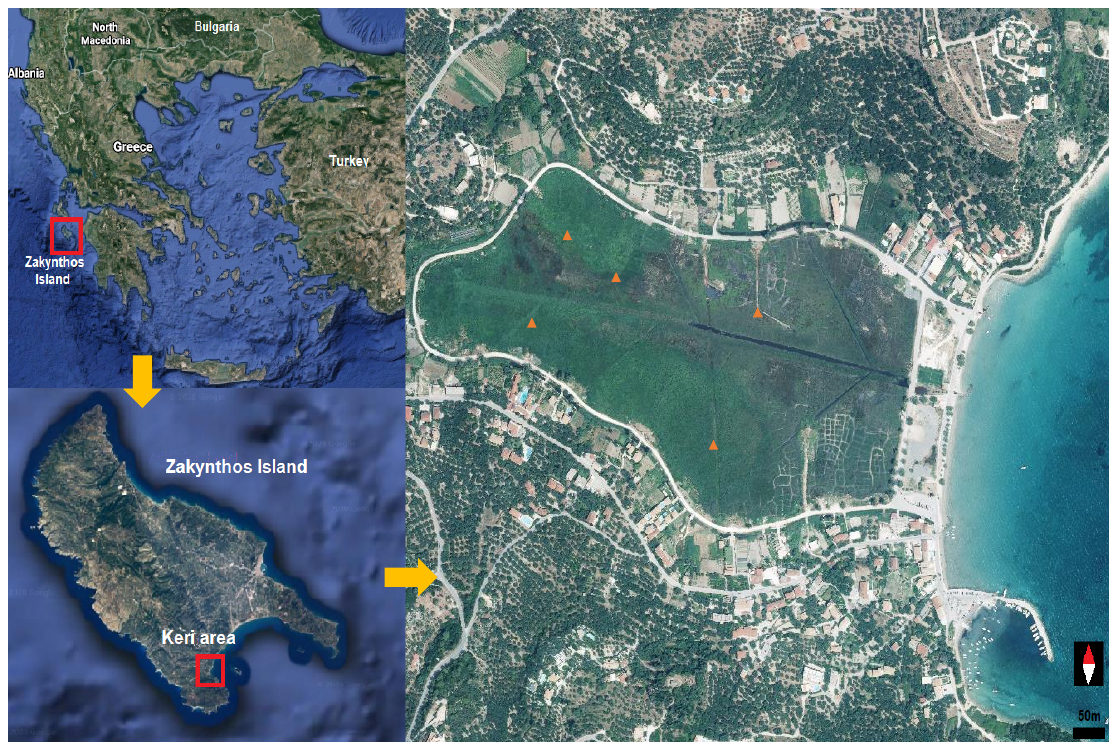

**Supplementary Figure 1.** The Keri Lake, at Zakynthos island, western Greece. The sampling sites are indicated with orange triangles.
