## Supplemental Figure2 for "Microbial bioprospecting for lignocellulose degradation at a unique Greek environment" (from another version of this preprint: version 1, 2020-08-29)

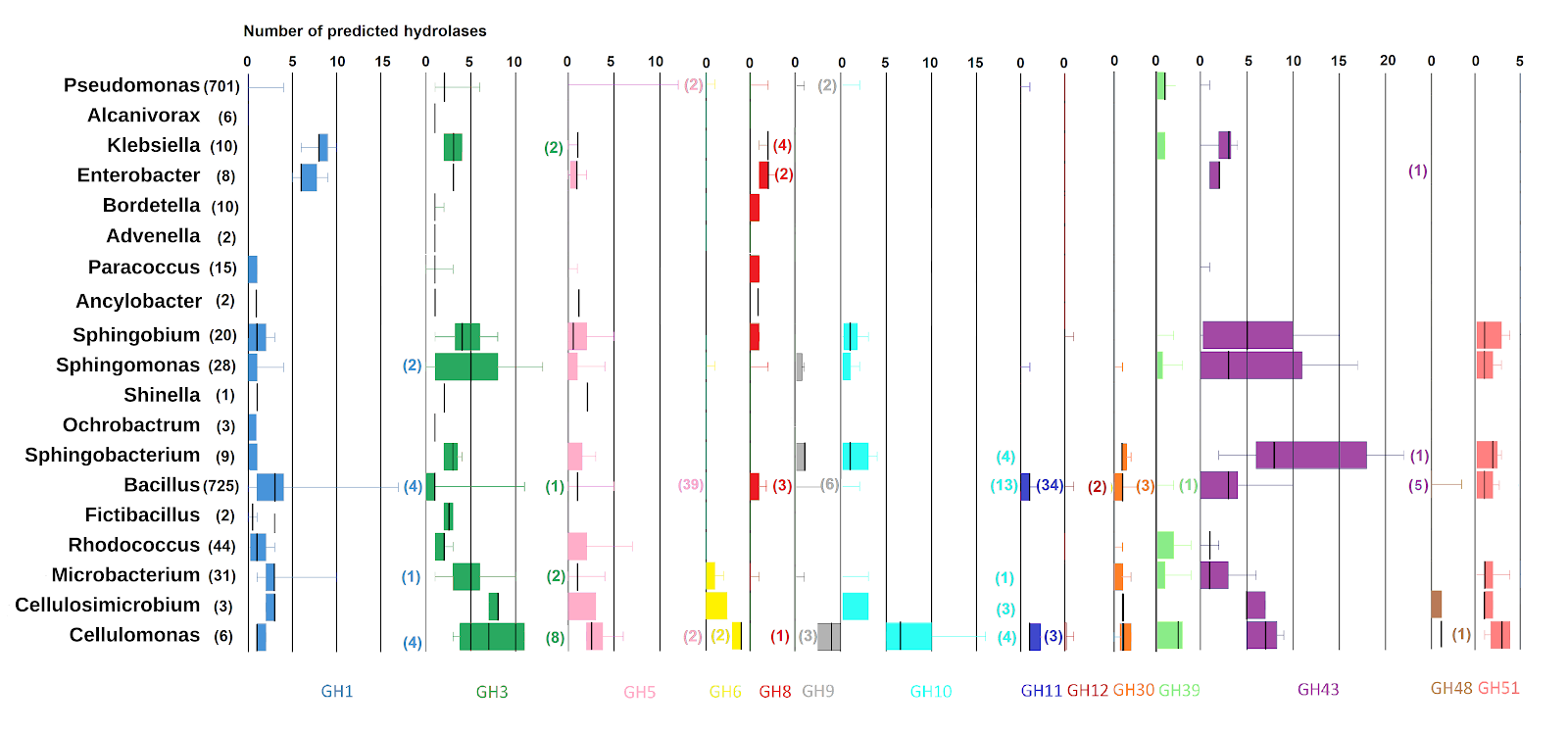

**Figure S2.** Predicted number of glycoside hydrolase genes involved in cellulose and xylan deconstruction, in available genomes, according to CAZy database. Colored boxes represent 25^th^ to 75^th^ percentiles, black horizontal lines represent medians, and error bars represent minimum and maximum values. Black brackets correspond to the number of sequenced genomes per genus and colored brackets to the number of biochemically characterized enzymes with activity corresponding to EC numbers 3.2.1.4, 3.2.1.21, 3.2.1.176, 3.2.1.91, 3.2.1.8 and 3.2.1.37. Enzymes belong primarily to the following families: Endoglucanases: GH5-GH9, GH12, GH44, GH45, GH48 and GH51, β-glucosidases: GH1, GH3, GH5, GH9 and GH30, Exoglucanases: GH5, GH6, GH9 and GH48, Endo-β-1,4-xylanases: GH5, GH8, GH10, GH11 and GH43, β-xylosidases: GH1, GH3, GH5, GH10, GH30, GH39, GH43 and GH51.
