## Supplemental Table1 for "Microbial bioprospecting for lignocellulose degradation at a unique Greek environment" (from another version of this preprint: version 1, 2020-08-29)

**Table S1**. Taxonomic identification of Keri isolates, based on 16S rRNA sequences. The highest score candidates from GenBank database and the highest similarity candidates from EzBioCloud database are shown.

| **Isolate** | **ATHUBA Culture Collection Acc. No.** | **16S rRna  GenBank Acc. No.** | **GenBank** | | **EzBioCloud** | |
| --- | --- | --- | --- | --- | --- | --- |
|  |  |  | **Strain** | **Identity (%)** | **Strain** | **Identity (%)** |
| ZKA1 | ATHUBa1401 | MT683172 | *Achromobacter pulmonis* R-16442 | 100 | *Achromobacter pulmonis* LMG 26696 | 99.85 |
| ZKA2 | ATHUBa1402 | MT683173 | *Pseudomonas aeruginosa* DSM 50071 | 100 | *Pseudomonas aeruginosa* JCM 5962 | 100 |
| ZKA3 | ATHUBa1403 | MT683174 | *Pseudomonas plecoglossicida* NBRC 103162 | 99 | *Pseudomonas putida* KT2440 | 100 |
| ZKA4 | ATHUBa1404 | MT683175 | *Bordetella tumbae* T6713-1-3b | 99 | *Bordetella tumbae* T6713-1-3b | 99.62 |
| ZKA5 | ATHUBa1405 | MT683176 | *Pseudomonas pseudoalcaligenes* Stanier 63 | 99 | *Pseudomonas oleovorans* subsp. oleovorans DSM 1045 | 98.73 |
| ZKA6 | ATHUBa1406 | MT683177 | *Pseudomonas plecoglossicida* NBRC 103162 | 99 | *Pseudomonas* sp. R17(2017) | 100 |
| ZKA7 | ATHUBa1407 | MT683178 | *Pseudomonas kilonensis* 520-20 | 99 | *Pseudomonas fluorescens* 2P24 | 99.86 |
| ZKA8 | ATHUBa1408 | MT683179 | *Achromobacter pulmonis* R-16442 | 100 | *Achromobacter pulmonis* LMG 26696 | 99.86 |
| ZKA9 | ATHUBa1409 | MT683180 | *Pseudomonas resinovorans* ATCC 14235 | 100 | *Pseudomonas resinovorans* LMG 26696 | 99.15 |
| ZKA10 | ATHUBa1410 | MT683181 | *Pseudomonas plecoglossicida* NBRC 103162 | 100 | *Pseudomonas* sp. R17(2017) | 100 |
| ZKA11 | ATHUBa1411 | MT683182 | *Pseudomonas oryzihabitans* L-1 | 99 | *Pseudomonas capeferrum* WCS358 | 99.93 |
| ZKA12 | ATHUBa1412 | MT683183 | *Pseudomonas plecoglossicida* NBRC 103162 | 99 | *Pseudomonas* sp. R17(2017) | 100 |
| ZKA13 | ATHUBa1413 | MT683184 | *Pseudomonas oryzihabitans* L-1 | 99 | *Pseudomonas capeferrum* WCS358 | 99.93 |
| ZKA14 | ATHUBa1414 | MT683185 | *Sphingobium yanoikuyae* NBRC 15102 | 99 | *Sphingobium yanoikuyae* ATCC 51230 | 99.78 |
| ZKA15 | ATHUBa1415 | MT683186 | *Microbacterium ginsengiterrae* DCY37 | 98 | *Microbacterium shaanxiense* CCNWSP60(T) | 98.34 |
| ZKA16 | ATHUBa1416 | MT683187 | *Paracandidimonas soli* IMT-305 | 97 | *Paracandidimonas soli* IMT-305 | 98.43 |
| ZKA17 | ATHUBa1417 | MT683188 | *Cellulosimicrobium cellulans* DSM 43879 | 99 | *Cellulosimicrobium funkei* ATCC BAA-886 | 99.64 |
| ZKA18 | ATHUBa1418 | MT683189 | *Paracandidimonas soli* IMT-305 | 98 | *Paracandidimonas soli* IMT-305 | 98.65 |
| ZKA19 | ATHUBa1419 | MT683190 | *Cellulomonas pakistanensis* NCCP-11 | 98 | *Cellulomonas hominis* JCM 12133 | 98.5 |
| ZKA20 | ATHUBa1420 | MT683191 | *Klebsiella aerogenes* NBRC 13534 | 99 | *Klebsiella aerogenes* KCTC 2190 | 99.86 |
| ZKA21 | ATHUBa1421 | MT683192 | *Microbacterium thalassium* DSM 12511 | 99 | *Microbacterium thalassium* IFO 16060 | 98.42 |
| ZKA22 | ATHUBa1422 | MT683193 | *Stenotrophomonas maltophilia* IAM 12423 | 99 | *Stenotrophomonas maltophilia* MTCC 434 | 99.72 |
| ZKA23 | ATHUBa1423 | MT683194 | *Shinella kummerowiae* CCBAU 25048 | 98 | *Sinorhizobium* sp. RAC02 | 98.46 |
| ZKA24 | ATHUBa1424 | MT683195 | *Ochrobactrum rhizosphaerae* PR17 | 99 | *Ochrobactrum pituitosum* CCUG 50899 | 99.71 |
| ZKA25 | ATHUBa1425 | MT683196 | *Sphingobium yanoikuyae* NBRC 15102 | 99 | *Sphingobium* sp. AP49 | 99.27 |
| ZKA26 | ATHUBa1426 | MT683197 | *Advenella kashmirensis* WT001 | 99 | *Advenella kashmirensis* subsp. *methylica* PK1 | 99.93 |
| ZKA27 | ATHUBa1427 | MT683198 | *Microbacterium sorbitolivorans* SZDIS-1-1 | 99 | *Microbacterium sorbitolivorans* SZDIS-1-1 | 99.86 |
| ZKA28 | ATHUBa1428 | MT683199 | *Pseudomonas composti* C2 | 100 | *Pseudomonas composti* CCUG 59231 | 100 |
| ZKA29 | ATHUBa1429 | MT683200 | *Paracoccus caeni* MJ17 | 99 | *Paracoccus caeni* MJ17 | 100 |
| ZKA30 | ATHUBa1430 | MT683201 | *Microbacterium pumilum* KV-488 | 99 | *Microbacterium pumilum* KV-488 | 99.21 |
| ZKA31 | ATHUBa1431 | MT683202 | *Advenella kashmirensis* WT001 | 99 | *Advenella kashmirensis* subsp. *methylica* PK1 | 99.93 |
| ZKA32 | ATHUBa1432 | MT683203 | *Pseudomonas plecoglossicida* NBRC 103162 | 99 | *Pseudomonas* sp. R17(2017) | 100 |
| ZKA33 | ATHUBa1433 | MT683204 | *Rhodococcus pyridinivorans* PDB9 | 100 | *Rhodococcus pyridinivorans* DSM 44555 | 99.86 |
| ZKA34 | ATHUBa1434 | MT683205 | *Parapedobacter pyrenivorans* P-4 | 99 | *Parapedobacter pyrenivorans* P-4 | 98.85 |
| ZKA35 | ATHUBa1435 | MT683206 | *Pseudomonas plecoglossicida* NBRC 103162 | 99 | *Pseudomonas* sp. R17(2017) | 100 |
| ZKA36 | ATHUBa1436 | MT683207 | *Pseudomonas aeruginosa* DSM 50071 | 99 | *Pseudomonas aeruginosa* JCM 5962 | 99.86 |
| ZKA37 | ATHUBa1437 | MT683208 | *Pseudomonas putida* NBRC 14164 | 99 | *Pseudomonas putida* ATH-43 | 99.86 |
| ZKA38 | ATHUBa1438 | MT683209 | *Achromobacter pulmonis* R-16442 | 100 | *Achromobacter pulmonis* LMG 26696 | 100 |
| ZKA39 | ATHUBa1439 | MT683210 | *Pseudomonas songnenensis* NEAU-ST5-5 | 98 | *Pseudomonas stutzeri* DSM 17088 | 99.78 |
| ZKA40 | ATHUBa1440 | MT683211 | *Stenotrophomonas pavanii strain LMG 25348* | 98 | *Stenotrophomonas indicatrix* WS40 | 100 |
| ZKA41 | ATHUBa1441 | MT683212 | *Pseudomonas plecoglossicida* NBRC 103162 | 99 | *Pseudomonas putida* KT2440 | 100 |
| ZKA42 | ATHUBa1442 | MT683213 | *Sphingomonas sanxanigenens* NX02 | 99 | *Sphingomonas sanxanigenens* DSM 19645 | 99.93 |
| ZKA43 | ATHUBa1443 | MT683214 | *Microbacterium resistens* DMMZ 1710 | 98 | *Microbacterium shaanxiense* CCNWSP60(T) | 98.34 |
| ZKA44 | ATHUBa1444 | MT683215 | *Sphingobacterium pakistanense* NCCP-246 | 98 | *Sphingobacterium pakistanense* NCCP-246 | 98.29 |
| ZKA45 | ATHUBa1445 | MT683216 | *Bordetella tumbae* T6713-1-3b | 99 | *Bordetella tumbae* T6713-1-3b | 99.65 |
| ZKA46 | ATHUBa1446 | MT683217 | *Microbacterium trichothecenolyticum* DSM 8608 | 99 | *Microbacterium trichothecenolyticum* DSM 8608 | 99.14 |
| ZKA47 | ATHUBa1447 | MT683218 | *Pseudomonas aeruginosa* DSM 50071 | 100 | *Pseudomonas aeruginosa* JCM 5962 | 100 |
| ZKA48 | ATHUBa1448 | MT683219 | *Cellulosimicrobium cellulans* DSM 43879 | 99 | *Cellulosimicrobium funkei* ATCC BAA-886 | 99.5 |
| ZKA49 | ATHUBa1449 | MT683220 | *Rhodococcus pyridinivorans* PDB9 | 99 | *Rhodococcus pyridinivorans* DSM 44555 | 99.86 |
| ZKA50 | ATHUBa1450 | MT683221 | *Pseudomonas pseudoalcaligenes* Stanier 63 | 99 | *Pseudomonas oleovorans* subsp. *Oleovorans* DSM 1045 | 98.73 |
| ZKA51 | ATHUBa1451 | MT683222 | *Pseudomonas* *pseudoalcaligenes* Stanier 63 | 99 | *Pseudomonas oleovorans* subsp. *Oleovorans* DSM 1045 | 98.8 |
| ZKA52 | ATHUBa1452 | MT683223 | *Alcanivorax dieselolei* B5 | 99 | *Alcanivorax dieselolei* B-5 | 99.86 |
| ZKA53 | ATHUBa1453 | MT683224 | *Ancylobacter pratisalsi* E130 | 99 | *Ancylobacter pratisalsi* E130 | 99.41 |
| ZKA54 | ATHUBa1454 | MT683225 | *Sinomicrobium oceani* SCSIO 03483 | 99 | *Sinomicrobium oceani* SCSIO 03483 | 100 |
| ZKA55 | ATHUBa1455 | MT683226 | *Pseudomonas knackmussii* B13 | 99 | *Pseudomonas kunmingensis* HL22-2 | 99.29 |
| ZKA56 | ATHUBa1456 | MT683227 | *Microbacterium esteraromaticum* DSM 8609 | 99 | *Microbacterium esteraromaticum* DSM 8609 | 99.64 |
| ZKA57 | ATHUBa1457 | MT683228 | *Microbacterium esteraromaticum* DSM 8609 | 99 | *Microbacterium esteraromaticum* DSM 8609 | 99.64 |
| ZKA58 | ATHUBa1458 | MT683229 | *Bacillus thuringiensis* ATCC 10792 | 100 | *Bacillus mobilis* 0711P9-1 | 100 |
| ZKA59 | ATHUBa1459 | MT683230 | *Fictibacillus nanhaiensis* JSM 082006 | 99 | *Fictibacillus nanhaiensis* JSM 082006 | 99.44 |
| ZKA60 | ATHUBa1460 | MT683231 | *Bacillus mojavensis* NBRC 15718 | 99 | *Bacillus halotolerans* ATCC 25096 | 99.93 |
| ZKA61 | ATHUBa1461 | MT683232 | *Bacillus thuringiensis* ATCC 10792 | 99 | *Bacillus mobilis* 0711P9-1 | 99.93 |
| ZKA62 | ATHUBa1462 | MT683233 | *Enterobacter tabaci* YIM Hb-3 | 99 | *Enterobacter tabaci* YIM Hb-3 | 99.86 |
| ZKA63 | ATHUBa1463 | MT683234 | *Sphingobacterium siyangense* SY1 | 99 | *Sphingobacterium siyangense* SY1 | 99.25 |
